## Supplemental Information for "The bile acid-sensitive ion channel is gated by Ca^2+^-dependent conformational changes in the transmembrane domain"

**Supplementary Information**

**Supplementary Fig. 1:** Function of hBASIC construct.

**Supplementary Fig. 2:** Biochemical and functional characterization of recombinant hBASIC.

**Supplementary Fig. 3:** Cryo-EM processing pipeline for 2 mM $Ca^{2+}$ data set.

**Supplementary Fig. 4:** 2mM $Ca^{2+}$ hBASIC cryo-EM map analysis.

**Supplementary Fig. 5:** Comparison of hBASIC to cASIC1a.

**Supplementary Fig. 6:** Cryo-EM processing workflow of EGTA data set.

**Supplementary Fig. 7:** EGTA, closed, hBASIC cryo-EM map analysis.

**Supplementary Figure 8.** EGTA, expanded, hBASIC cryo-EM map analysis.

**Supplementary Figure 9.** Example traces and statistical analysis of $IC_{50}$.

**Supplementary Figure 10.** Current-voltage and conductance-voltage analysis.

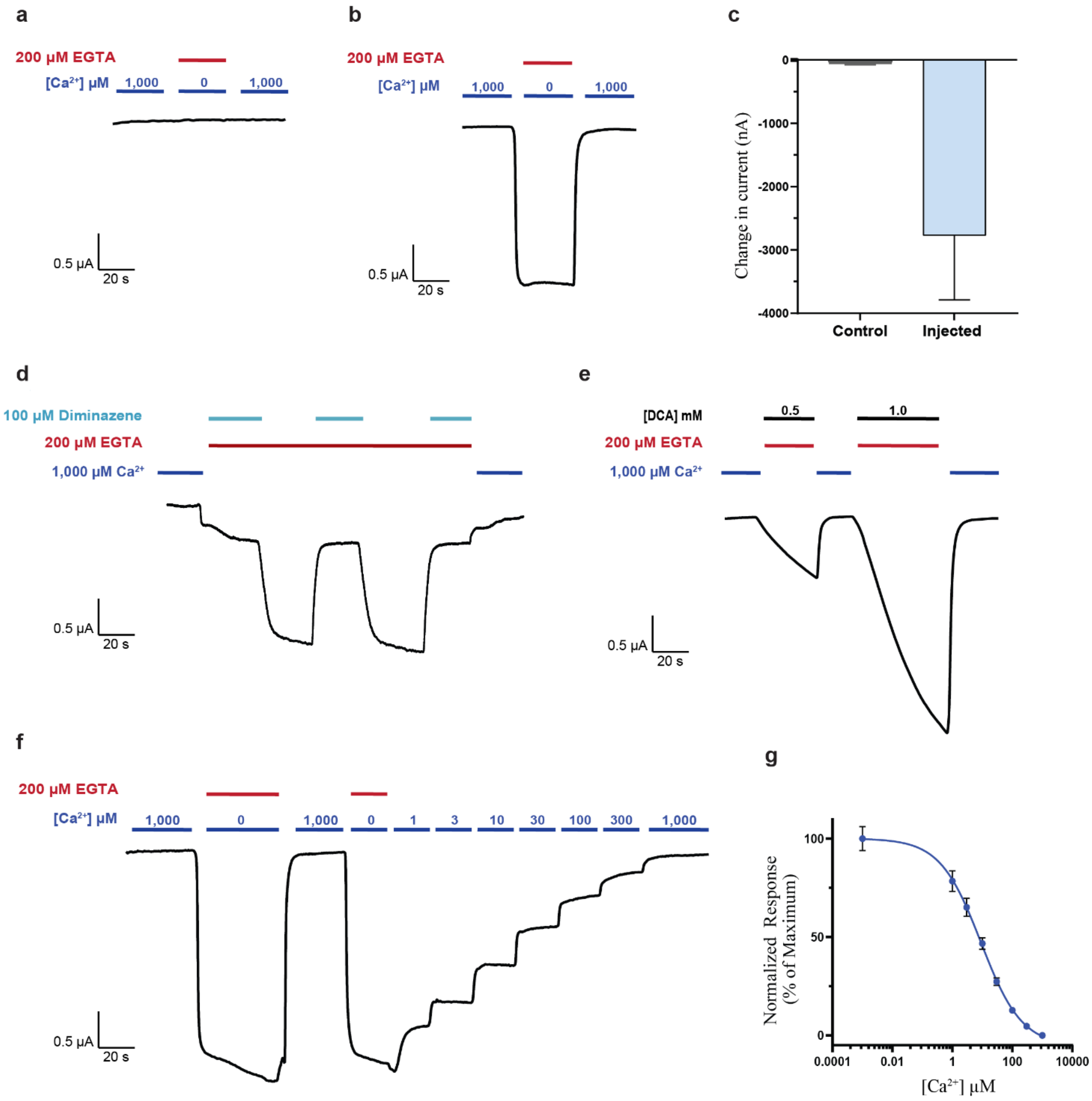


**Supplementary Fig. 1. Function of hBASIC construct. (a-b)** Example recording of **(a)** oocyte control and **(b)** oocyte expressing hBASIC when recorded in the presence of 1 mM $Ca^{2+}$ or 0.2 mM EGTA. **(c)** Change in current (nA) upon $Ca^{2+}$ removal from the bath between control oocytes and hBASIC injected oocytes. **(d)** hBASIC's inhibitor, diminazene, attenuates $Ca^{2+}$ chelation activated currents in oocytes expressing hBASIC. **(e)** $Ca^{2+}$ chelation activated currents can be potentiated by increasing amounts of the bile acid, deoxycholic acid (DCA). **(f-g)** There is no significant difference in potency of $Ca^{2+}$ inhibition of hBASIC by $Ca^{2+}$ when perfusion of [$Ca^{2+}$] is performed in a 'reverse order' (i.e. increasing [$Ca^{2+}$], instead of decreasing [$Ca^{2+}$]). **(f)** A representative trace of the control experiment with the corresponding statistics analysis **(g)** that show the $IC_{50}$ is 8µM ± 2.

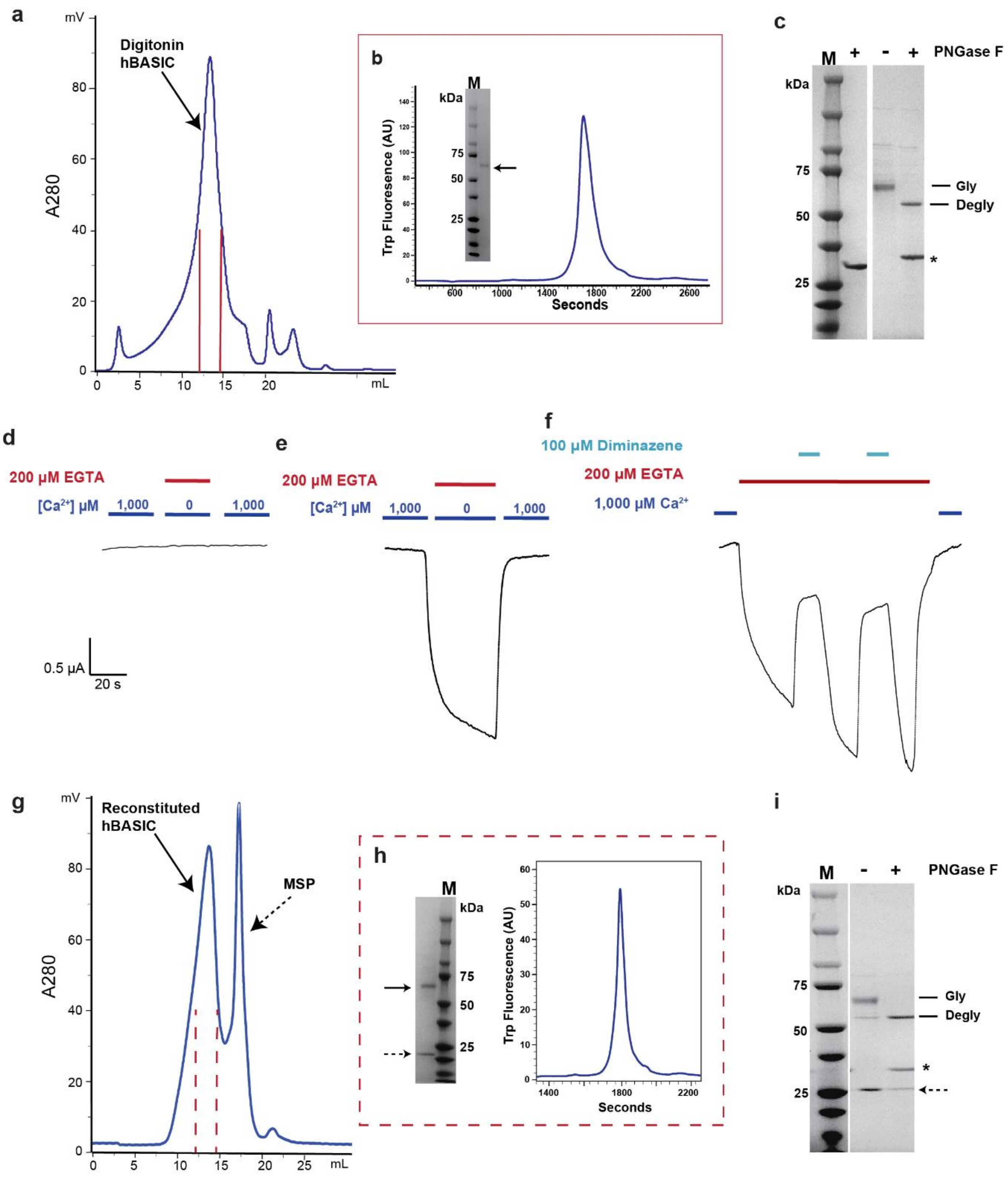


**Supplementary Fig. 2. Biochemical and functional characterization of recombinant hBASIC. (a)** Size exclusion chromatography (SEC) chromatogram of digitonin extracted hBASIC expressed in HEK293 cells. Solid red lines indicate peak fraction corresponding to hBASIC. **(b)** Fluorescent size exclusion chromatography (FSEC) and SDS-PAGE gel analysis of digitonin hBASIC sample from peak fraction of

SEC elution. **(c)** SDS-PAGE analysis of digitonin solubilized hBASIC with and without PNGase F treatment. Lines indicate glycosylated and deglycosylated hBASIC. Star indicates PNGase F. **(d-f)** Representative TEVC current trace recordings of **(d)** uninjected oocytes and **(e-f)** oocytes injected with digitonin purified hBASIC reconstituted into liposomes. Injected oocytes demonstrate inward sodium currents in response to the removal of $Ca^{2+}$ via EGTA chelation. **(f)** The EGTA chelated currents are attenuated with hBASIC pore blocker, diminazene. **(g)** Example SEC chromatogram of nanodisc reconstituted hBASIC. Solid arrow represents nanodisc embedded hBASIC, while a dashed arrow represents membrane scaffold protein (MSP). Elution fraction indicated with dashed red lines. **(h)** SDS-page and FSEC analysis of nanodisc embedded hBASIC from SEC elution. **(i)** SDS-PAGE analysis of nanodisc-embedded hBASIC with and without PNGase F treatment. Lines indicate glycosylated and deglycosylated hBASIC. Star indicates PNGase F. Dashed arrow indicates MSP protein.

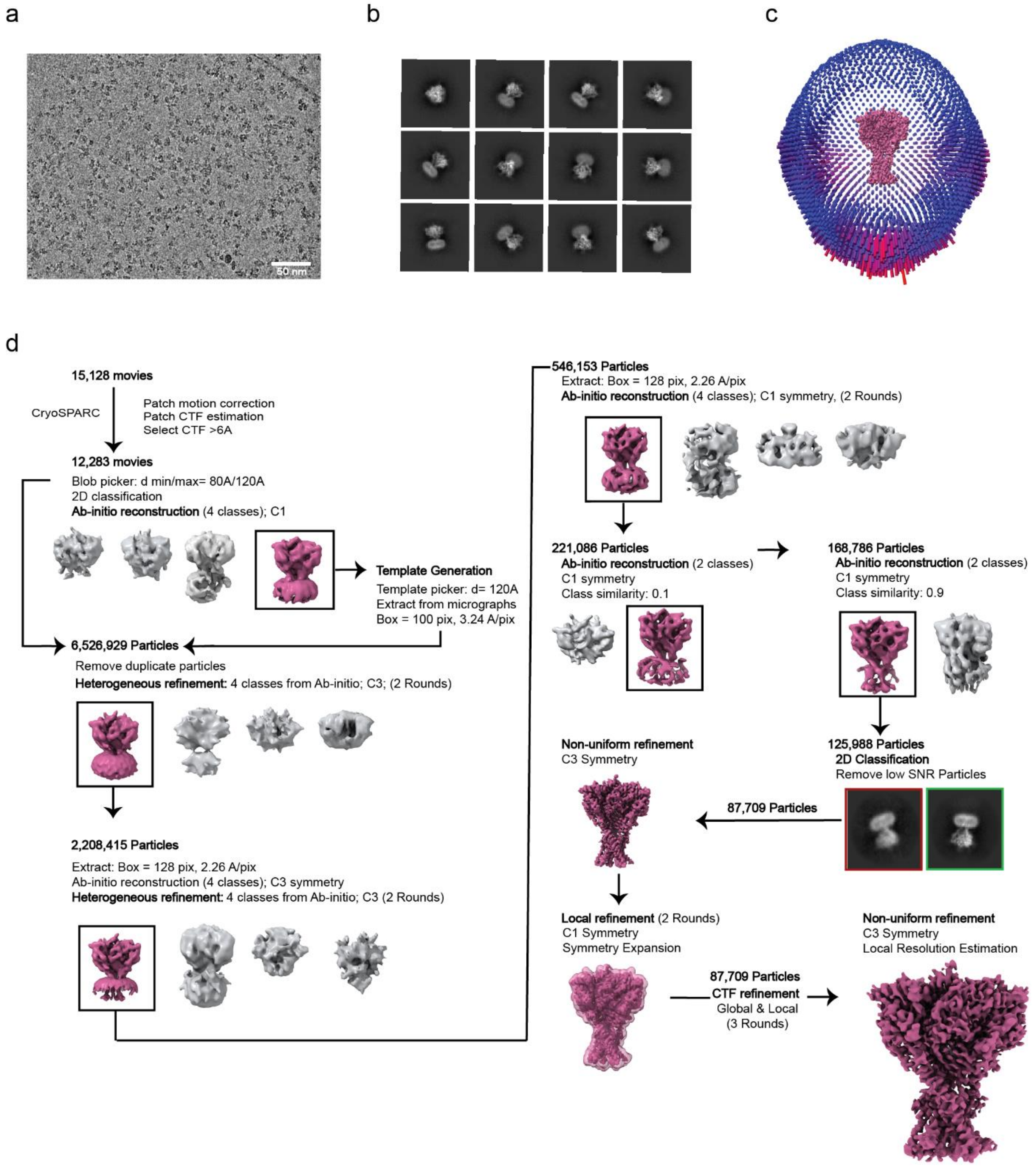


**Supplementary Fig. 3. Cryo-EM processing pipeline for 2 mM $Ca^{2+}$ data set. (a)** Representative micrograph from data set. **(b)** Selected 2D classes and **(c)** angular distribution of particles included in the final map. **(d)** Data processing workflow.

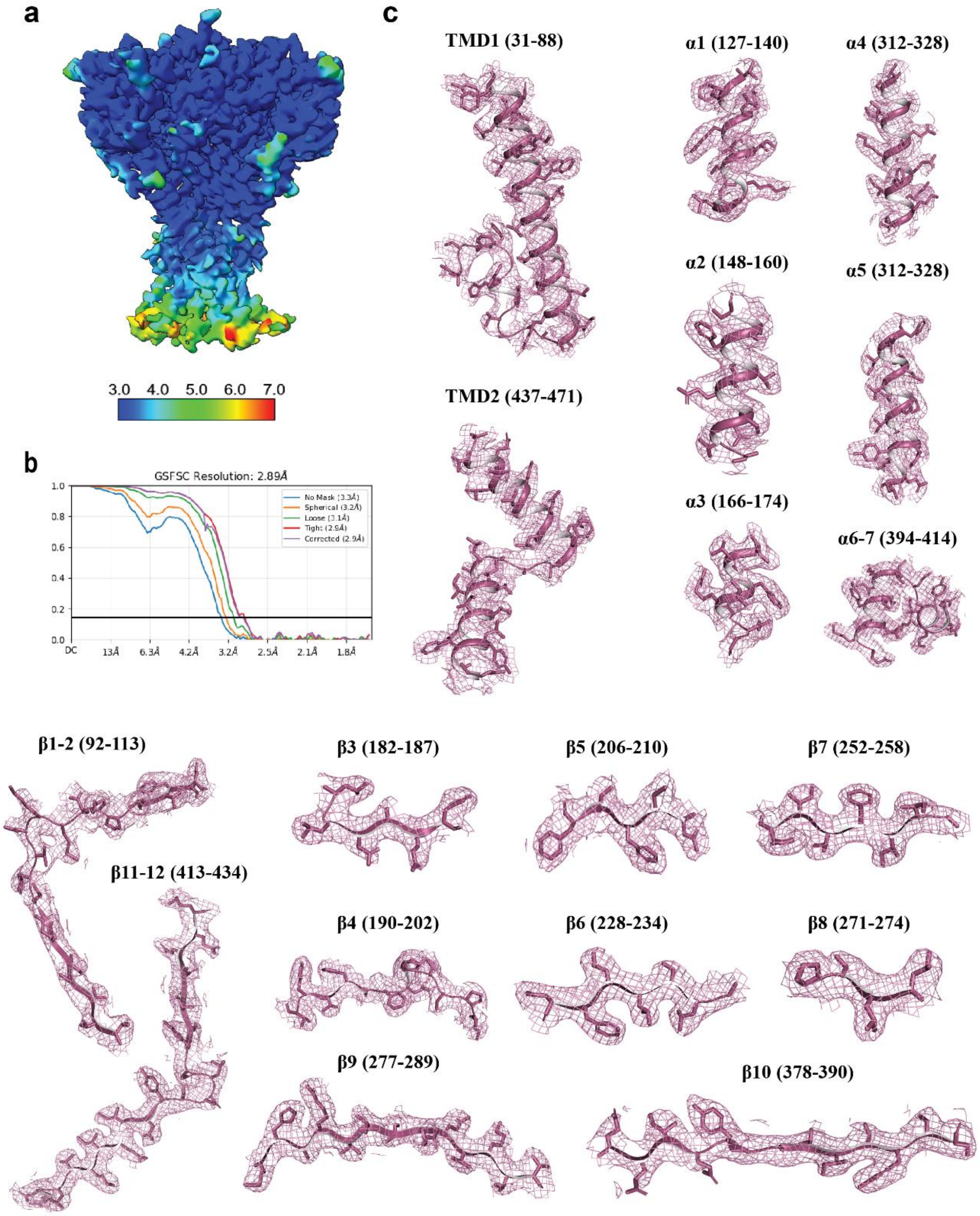


**Supplementary Fig. 4. 2mM $Ca^{2+}$ hBASIC cryo-EM map analysis. (a)** Local resolution and **(b)** Fourier shell correlation (FSC) plot of 2mM $Ca^{2+}$ hBASIC map. **(c)** Density associated with TMDs, α-helices, and β-sheets. Isomesh map features are contoured at 6.0 σ and within 2Å associated with each feature.

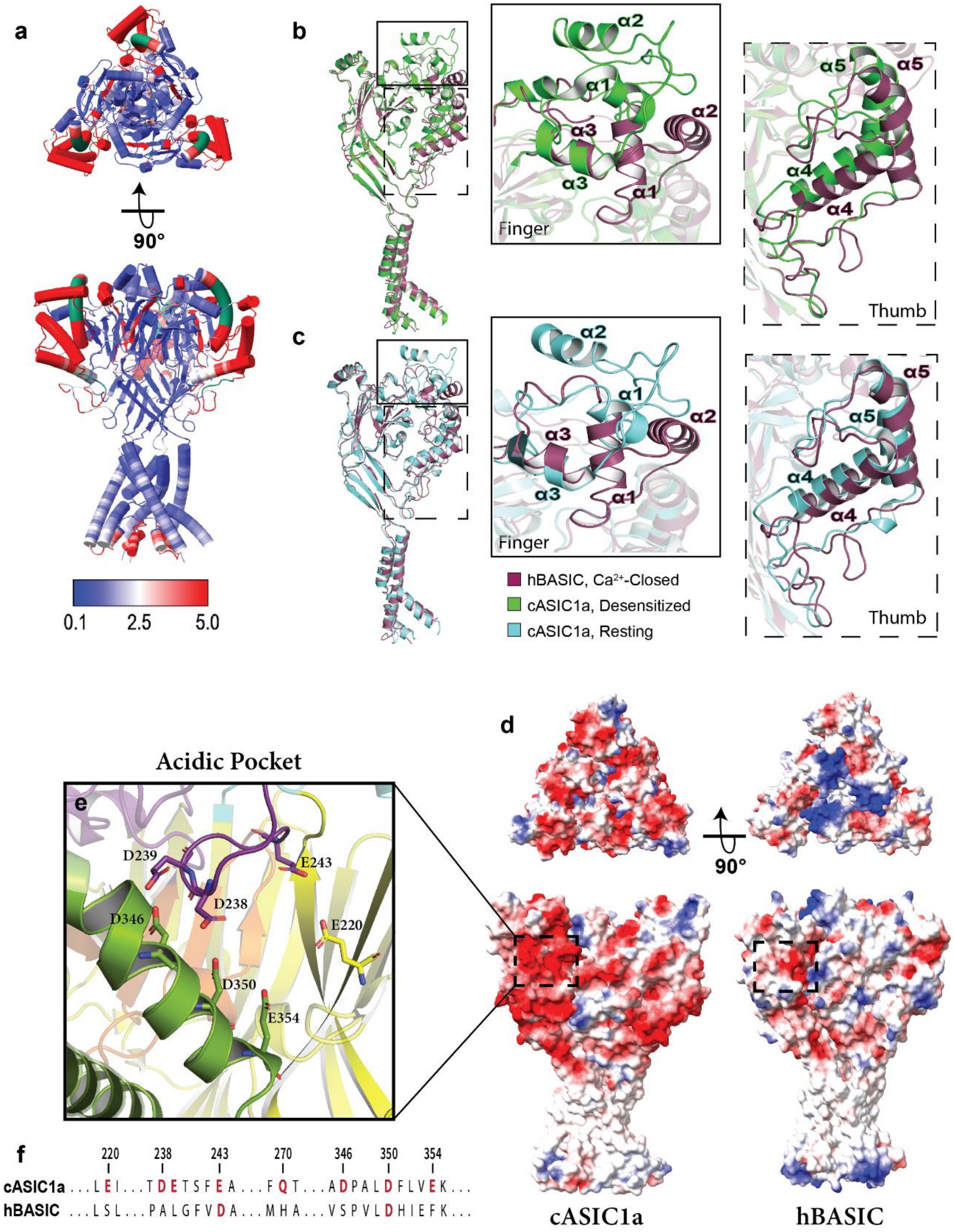


**Supplementary Fig. 5. Comparison of hBASIC to cASIC1a. (a)** rmsd of $Ca^{2+}$ hBASIC in comparison to cASIC1a, desensitized [PDB:6VTK]. Green represents unique inserts, Blue = high conservation, Red = low conservation. **(b-c)** hBASIC with $Ca^{2+}$ map superimposed onto cASIC1a desensitized [PDB:6VTK] **(b)** and resting [PDB: 6VTL] **(c)**, highlighting regions of low conservation, the finger and thumb. **(d)** Models of cASIC1a and hBASIC colored by electrostatic potential, with red representing regions of negative potential through blue for positive potential. A dashed box indicates the acidic pocket of cASIC1a and the equivalent region in hBASIC. **(e)** The acidic pocket of cASIC1a, a pocket formed by intersecting regions of the finger, thumb, and palm domain. **(f)** Sequence comparison of the acidic pocket between cASIC1a and hBASIC, showing lack of sequence conservation of these acidic residues.

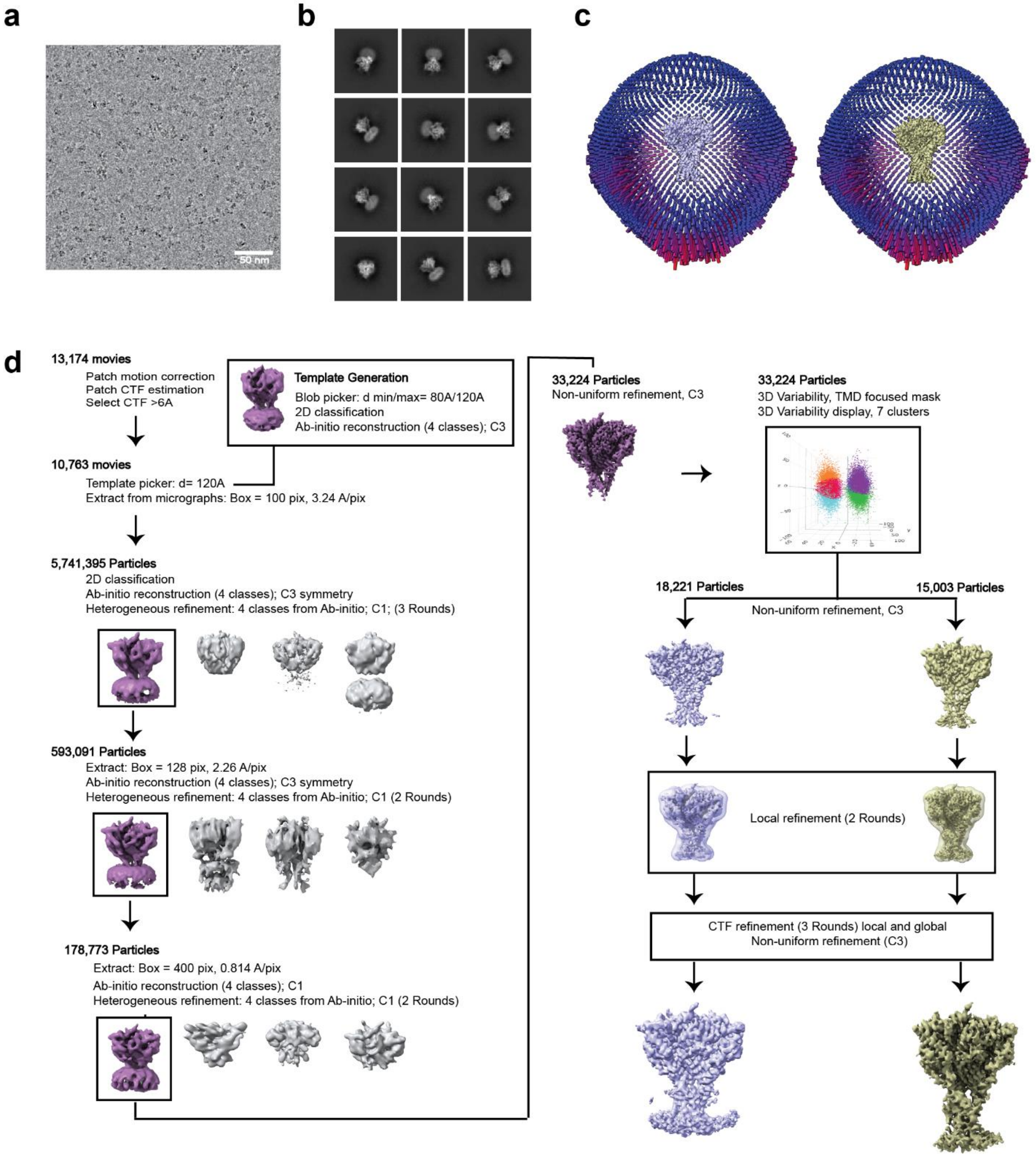


**Supplementary Fig. 6. Cryo-EM processing workflow of EGTA data set. (a)** Representative micrograph from data set. **(b)** Selected 2D classes and **(c)** angular distribution of particles included in the final maps. **(d)** Data processing workflow.

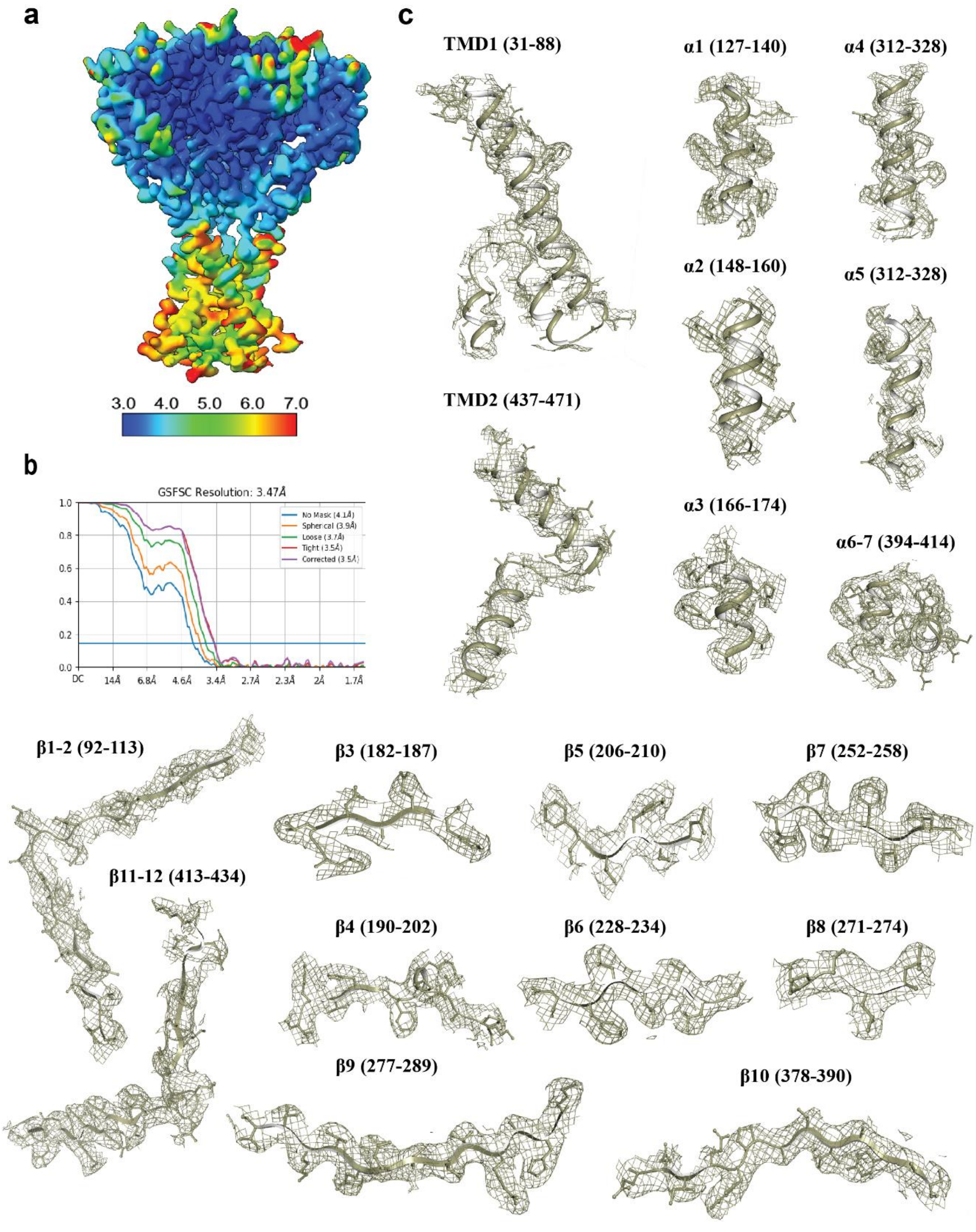


**Supplementary Fig. 7. EGTA, closed, hBASIC cryo-EM map analysis. (a)** Local resolution and **(b)** Fourier shell correlation (FSC) plot of EGTA, closed, hBASIC map. **(c)** Density associated with TMDs, α-helices, and β-sheets. Isomesh map features are contoured at 4.0 σ and within 2Å associated with each feature.

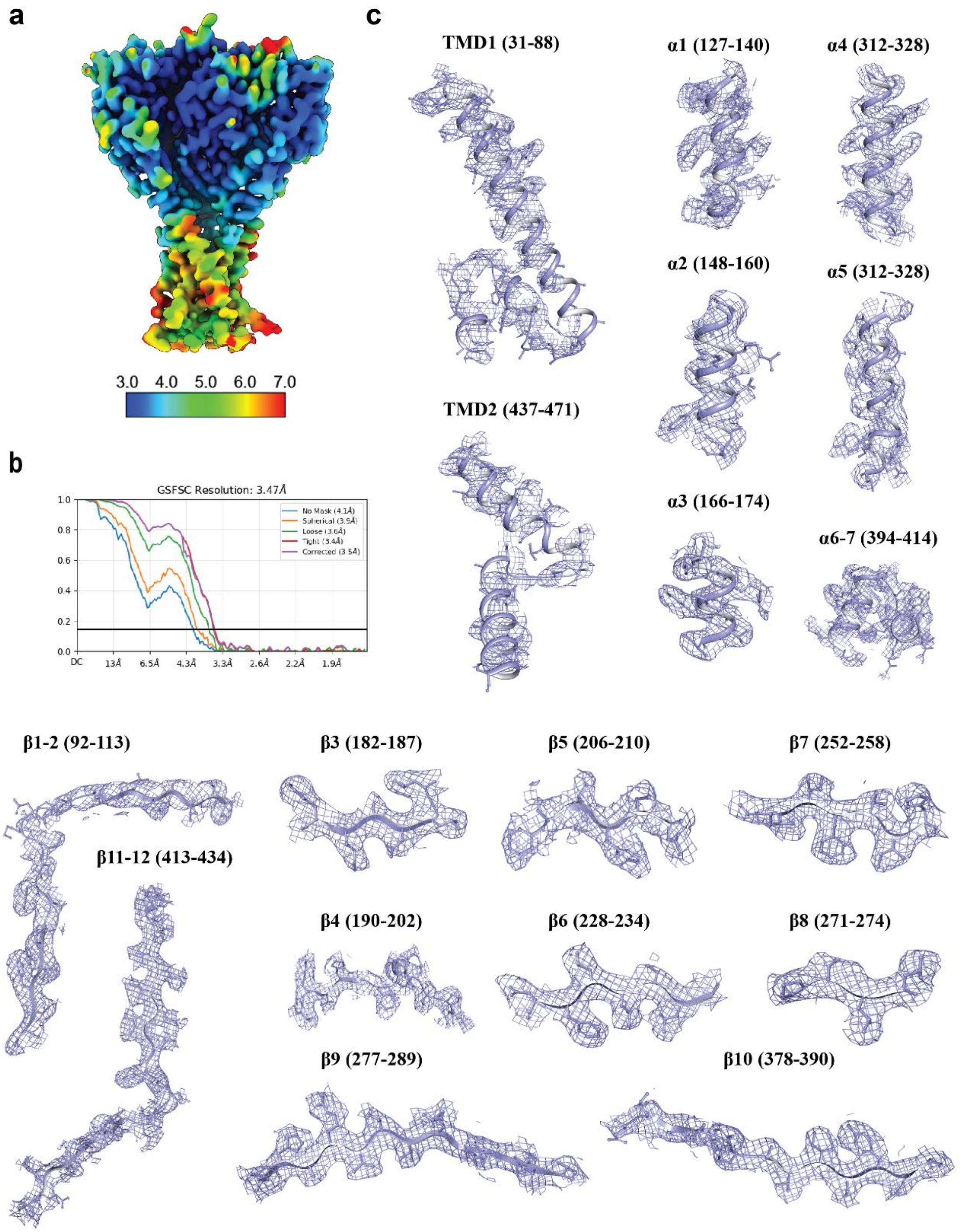


**Supplementary Fig. 8. EGTA, expanded, hBASIC cryo-EM map analysis.** **(a)** Local resolution estimation of EGTA, expanded map. **(b)** Fourier shell correlation (FSC) plot of corresponding map. **(c)** Density, contoured at 4.0 σ and within 2Å associated with TMDs, α-helices, and β-sheets.

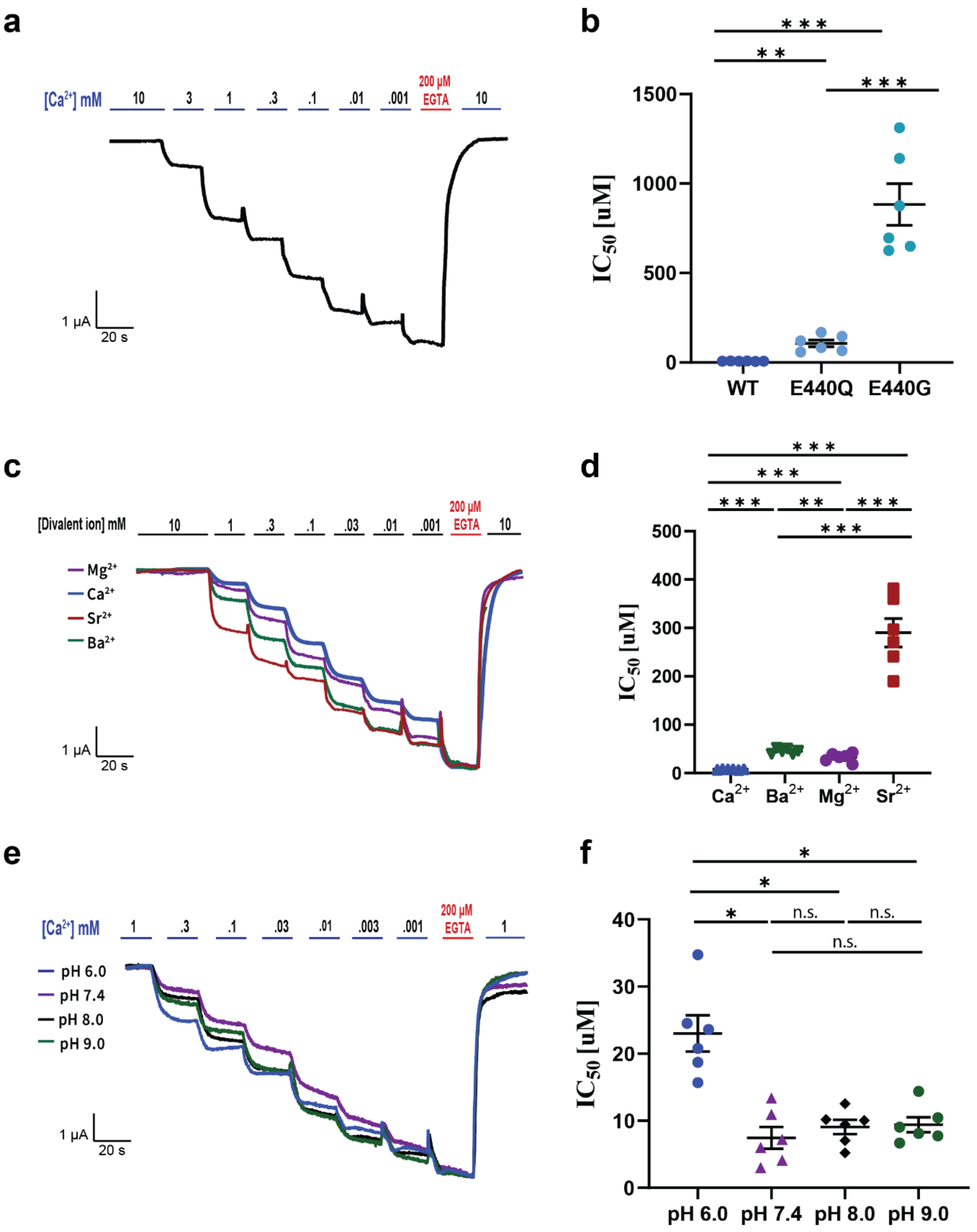


**Supplementary Fig. 9. Example traces and statistical analysis of $IC_{50}$. (a)** Example recording of an inhibition dose-response curve of $Ca^{2+}$ of an oocyte injected with mutant E440Q. **(b)** Statistical analysis of

$IC_{50}$ of WT, and mutants, E440Q and E440G. WT versus E440Q, P= 1.4 E-4; WT versus E440G, P=9.9 E-6, E440Q versus E440G, P=3.1E-5. **(c)** Example traces of dose-response recordings of oocytes expressing hBASIC for inhibition by $Mg^{2+}$, $Ca^{2+}$, $Sr^{2+}$, and $Br^{2+}$. **(d)** Statistical analysis of $IC_{50}$ of inhibition by divalent ions. $Ca^{2+}$ versus $Ba^{2+}$, P= 1.1 E-7; $Ca^{2+}$ versus $Mg^{2+}$, P=2.7 E-5, $Ca^{2+}$ versus $Sr^{2+}$, P=1.2 E-6, $Ba^{2+}$ versus $Mg^{2+}$, P=4.7 E-4, $Ba^{2+}$ versus $Sr^{2+}$, P=7.3 E-6, $Mg^{2+}$ versus $Sr^{2+}$, P=2.9 E-6. **(e)** Example traces of dose-response recordings of oocytes expressing hBASIC for inhibition by $Ca^{2+}$ at pH 6.0. **(f)** Statistical analysis of $IC_{50}$ of inhibition by $Ca^{2+}$ at different pH values. pH6 versus pH7.4, P= 0.019; pH6 versus pH8, P=0.026, pH6 versus pH9, P=0.028, pH7.4 versus pH8, P=0.127, pH7.4 versus pH9, P=0.103, pH8 versus pH9, P=0.42. **(b, d, f)** Statistical significances are shown as $*P < 0.05$, $**P < 0.01$, $***P < 0.001$. Significances were evaluated using two-tailed Student's *t*-tests.

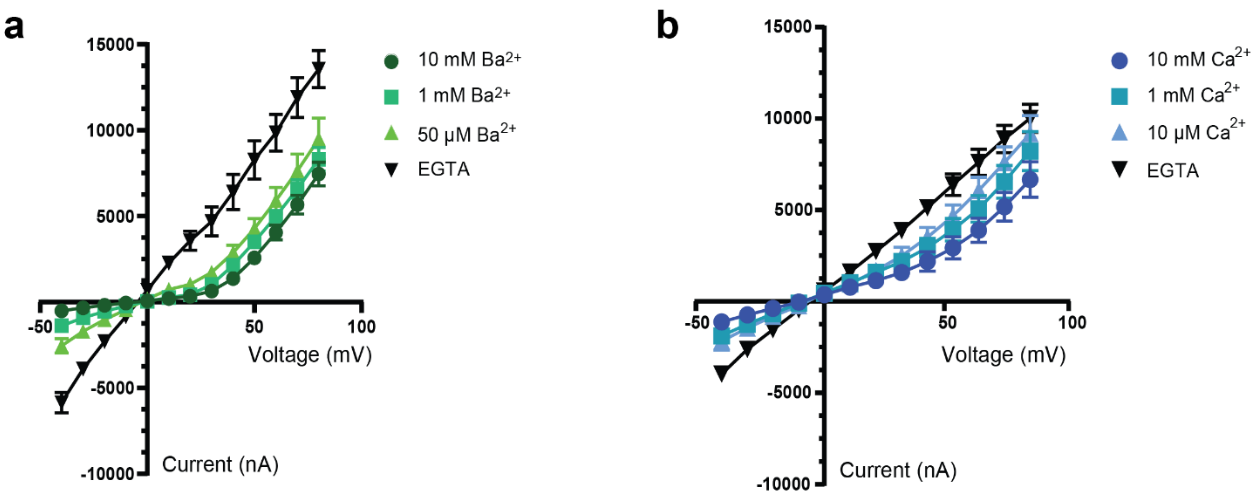


**Supplementary Fig. 10. Current-voltage and conductance-voltage analysis. (a)** Current-voltage (IV) plot of oocytes (n=7) injected with wild-type hBASIC, recording various amounts of $Br^{2+}$. **(b)** IV plot of oocytes (n=7) injected with mutant hBASIC, E440Q, recording various amounts of $Ca^{2+}$.
